# Can quantifying morphology and TMEM119 expression distinguish between microglia and infiltrating macrophages after ischemic stroke and reperfusion in male and female mice?

**DOI:** 10.1101/2020.09.23.310433

**Authors:** Kimberly F. Young, Rebeca Gardner, Victoria Sariana, Susan A. Whitman, Mitchell J. Bartlett, Torsten Falk, Helena W. Morrison

## Abstract

**Background:** Ischemic stroke is an acquired brain injury with gender dependent outcomes. A persistent obstacle in understanding the sex-specific neuroinflammatory contributions to ischemic brain injury is distinguishing between resident microglia versus infiltrating macrophages—both phagocytes—and determining cell population specific contributions to injury evolution and recovery processes. Our purpose was to identify microglial and macrophage populations regulated by ischemic stroke using morphology analysis and the presence of microglia transmembrane protein 119 (TMEM119). Second, we examined sex and menopause differences in microglia/macrophage cell populations after an ischemic stroke.

**Methods:** Male and female, premenopausal and postmenopausal, mice underwent either 60-min of middle cerebral artery occlusion and 24-h of reperfusion or sham surgery. The accelerated ovarian failure model was used to model post-menopause. Brain tissue was collected to quantify infarct area and for immunohistochemistry and western blot methods. Ionized calcium-binding adapter molecule, TMEM119, and confocal microscopy were used to analyze microglia morphology and TMEM119 area in ipsilateral brain regions. Western blot was used to quantify protein quantity.

**Results:** Post-stroke injury is increased in male and female post-menopause mice versus pre-menopause female mice (p<0.05) with differences primarily occurring in caudal sections. After stroke, microglia underwent a region, but not sex group, dependent transformation into less ramified cells (p<0.0001). However, the number of phagocytic microglia were increased in distal ipsilateral regions of postmenopausal mice versus the other sex groups (p<0.05). The number of TMEM119 positive cells was decreased in proximity to the infarct (p<0.0001) but without a sex group effect. Two key findings prevented distinguishing microglia from systemic macrophages. First, morphological data were not congruent with TMEM119 immunofluorescence data. Cells with severely decreased TMEM119 immunofluorescence were ramified, a distinguishing microglia characteristic. Second, whereas TMEM119 immunofluorescence area decreased in proximity to the infarcted area, TMEM119 protein quantity was unchanged in ipsilateral hemisphere regions using western blot methods.

**Conclusions:** Our findings suggest that TMEM119 is not a stable microglia marker in male and female mice in the context of ischemic stroke. Until TMEM119 function in the brain is elucidated, its use to distinguish between cell populations following brain injury with cell infiltration is cautioned.

## INTRODUCTION

Ischemic stroke is a significant cause of mortality and morbidity in the United States, with notable gender differences in patient outcomes [1]. In humans, the average age of ischemic stroke is 74 in men and 76 in women, with women experiencing an increase in stroke severity and incidence post-menopause. As such, biological sex and menopause status are important variables to consider in stroke studies that investigate gender-based differences [1-4]. Post-stroke neuroinflammation includes a complex, multi-faceted, and often sex specific cellular response to injury that contributes to brain recovery and repair after ischemic stroke [5-11]. However, in excess, neuroinflammatory responses can exacerbate injury in the acute phase [12] and may contribute to conditions such as cognitive decline [13] and Alzheimer’s disease [14]. Microglia are the resident brain immune cells and a key player in the neuroinflammatory response following an ischemic stroke [8, 15, 16]. Infiltrating macrophages/neutrophils also contribute to inflammatory responses in the brain; however, the timing of their arrival, numbers, and distinct functions are highly debated [15, 17-21]. Moreover, due to their common lineage [22, 23], it is difficult to distinguish microglia from infiltrating macrophages/neutrophils, further complicating our understanding of cell specific neuroinflammatory responses. This limitation impairs our ability to provide therapies that precisely target key contributors to injury resolution versus exacerbation in male and female stroke patients.

Morphology, parenchymal distribution, and a unique transcriptional profile are key distinguishing characteristics between these two similar cell populations [23-26]. Microglia are highly ramified cells that become less ramified in proximity to an injury [24], whereas macrophages are consistently amoeboid and have yet to be described as ramified cells. Identified through transcription data, transmembrane protein 119 (TMEM119) is a protein specific to microglia although its function and involvement in the brain’s injury response is yet unknown [27-29]. Its usefulness as a unique microglia protein marker was investigated and shown to have stable and specific protein expression in microglia in response to lipopolysaccharide (LPS) induced neuroinflammation and optic nerve crush [30]. However, its application in preclinical models of ischemia is limited [31]. Combining two *in-situ* methodologies, morphological analysis [32] and TMEM119 immunofluorescence and protein expression [30], our purpose was to definitively identify microglial and macrophage populations regulated by ischemic stroke. Our secondary purpose was to test for differences in sex and, in females, the effect of menopause on microglia/macrophage cell populations after an ischemic stroke. Utilizing a sixty-minute ischemia and twenty-four-hour reperfusion stroke protocol, we investigated if microglia transitioned into less ramified cells in proximity to the infarct similarly among our sex groups— male, pre-menopause and post-menopause mice. The TMEM119 immunofluorescence was also investigated among sex groups, to test how far the different cell populations might extend past the infarct border.

## METHODS

### Animals

All animal handling and experiments were performed according to methods approved by and in compliance with the University of Arizona Institutional Animal Care and Use Committee and according to the National Institutes of Health guide for the care and use of laboratory animals (protocol approval number 14-539). All animals were housed in rooms with a 12-h light/dark schedule (7 am - 7 pm) with food and water available *ad libitum*. Male and female C57Bl6/J mice were purchased from Jackson Laboratories (Bar Harbor, ME). The experimental unit for these studies is a single mouse and all sample sizes, along with statistical analyses used, are reported within the results and/or figures. A surgical control, sham, was used to compare microglial morphology and TMEM119 to healthy tissue while accounting for the general surgical procedure (described below). All mice were randomly assigned to either the sham or stroke condition. The ARRIVE guidelines were used for transparent reporting of research methods and findings [33].

### Postmenopause model

The accelerated ovarian failure model was used to model human post-menopause in this study of ischemic stroke [34, 35]. Similar to previous publications [34, 36], 5 wk old female mice were injected for 21 days with 4-vinylcyclohexene diepoxide (VCD, 160 mg/kg/*i*.*p*./day, Millipore Sigma #94956, St. Louis, MO). Follicle depletion and ovarian cessation was assessed ∼65 days after first injection via vaginal lavage, and ovarian failure was confirmed by observing 15 days of persistent diestrus prior to tissue collection. Because this is among the first applications of the post-menopause model in ischemic stroke research, we also tested the effect of the VCD injections themselves to exacerbate brain injury after ischemic stroke in male mice. Male mice (C57Bl6/J, 5 weeks) were treated with VCD for 21 days (160 mg/kg/*i*.*p*./day) followed by regular animal care until the stroke procedure.

### Ischemic stroke

A transient ischemic stroke was delivered using the filament method as previously published [24, 37]. Ischemic stroke was induced by temporary occlusion of the right middle cerebral artery (MCA) in anesthetized 16-week old mice (1–2% isoflurane in a 0.4 L/min medical air/0.1 L/min oxygen mixture). A filament was advanced, via the internal common carotid artery, to the ostea of the middle cerebral artery. The ischemic period for all experiments was 60 min, continuously verified by laser Doppler measures of relative cerebral blood flow to the MCA territory (Perimed Periflux 5000, North Royalton, OH). The sham procedure included all elements up to filament placement. Following 24 h of reperfusion, animals were exsanguinated and perfused with 0.01M phosphate buffered saline. For immunohistochemistry (IHC), brain tissue was removed and fixed in 4% paraformaldehyde for 24 hours followed by a 30% sucrose solution for 72 hours. For western blot methods, brain tissue was removed and rapidly frozen using 2-methylbutane (Thermo Fisher Scientific, Cat# O3551-4) at -50°C to prevent tissue cracking. All tissue was stored at -80°C until use.

### Immunohistochemistry

Fixed tissue was sectioned into 50 µm coronal sections (Leica cryostat CM1850, Buffalo Grove, IL) and stored at -20°C in a cryoprotectant solution (50% 50mM PBS, 30% ethylene glycol, 20% glycerol) until IHC experiments. A random and unbiased selection of tissue sections between Bregma 0 to +1 corresponding to ∼ 4 mm from the frontal pole was the basis for IHC methods and quantification of microglia morphology as well as the area of ionized calcium binding adaptor molecule 1 (IBA1) and TMEM119 immunofluorescence. Free floating brain sections were first blocked in 10% horse serum (Vector Laboratories, S-2000-20, Burlingame, CA) and buffer solution (0.01M PBS, 0.05% Triton, and 0.04% NaN_3_) for 1 h followed by a 72 h incubation with primary antibodies as appropriate: rabbit anti-IBA1 at 1:1000 (Wako, 019-19741, Madison, WI) and rat anti-TMEM119 at 1:250 (abcam 209064, Cambridge, UK) [24, 37]. The TMEM119 antibody has been validated for its specificity for microglia and its use in health and disease in an IHC preparation by others [25, 30]. In this case, healthy brain is considered a positive control, as well as a comparison group. A 4-hour incubation of 1:250 secondary antibodies (Jackson ImmunoResearch Laboratories, West Grove, PA) followed: donkey anti-rabbit Alexa 488 (711-546-152); donkey anti-rat Alexa 594, (712-585-150). All tissue was incubated in solutions common to all groups to avoid group/batch differences. All reactions were carried forward at room temperature; washes between incubations were with 0.01M PBS for 15 min. Vectashield (Vector Laboratories, H-1000) was used to coverslip mounted tissue.

### Imaging

Images were acquired on a confocal microscope (Zeiss NLO 880, San Diego, CA) in tissue regions in proximity to the infarct area. A 40X objective (236.16 × 236.16-micron area) was used for morphology analysis and a 20X objective (473.33 × 473.33-micron area) was used for percent area IBA1 and TMEM119 analysis. Z-stacks were compressed to a 2D image using maximum intensity, channels split and files saved as .tiff files using ImageJ software (NIH, v1 53c). Thresholding for the area of IBA1 and TMEM119 positive immunofluorescence was carried out using ImageJ and based on sham images; threshold parameters were consistent for all images. The percent area was recorded for all IBA1 and TMEM119. The number of cell somas were counted in the IBA1 channel, an approximation of the number of cells imaged in each frame, and all percent area data was divided by this cell count.

### Microglia Morphology Analysis

The extent of ramified microglial morphology was quantified using an objective and computer-aided skeleton analysis method as previously published in detail [32]. A series of ImageJ plugins (i.e. adjust brightness, unsharp mask, and despeckle) were temporarily applied in order to ensure adequate cell process visualization before the conversion to binary and skeletonized images. The skeletonized representations of original photomicrographs were used for data collection. The AnalyzeSkeleton (2D/3D) plugin (developed and maintained by Arganda-Carreras et al. [38]) was used to tag elements of microglia skeletons as processes (orange slab voxels) and endpoints (blue) for data collection. Microglia morphology was operationalized by reporting the number of microglia endpoints and process length. We summarized the number of endpoints and process length from AnalyzeSkeleton (2D/3D) plugin data output and all data were divided by cell soma counts, an approximation of the number of cells imaged in each frame. All analysis was carried out by researchers blinded to the sex group.

### Western Blot

In a different cohort of mice, tissue was collected after the ischemic stroke procedure, immediately frozen and stored until sectioning. Frozen tissue was sectioned into 2 mm coronal sections and a 1 mm diameter biopsy tool (Harris Micro-punch, Millipore-Sigma, Z708658) was used to collect tissue punches in ipsilateral regions distal and proximal to the infarcted tissue similar to IHC regions (**Figure 3B**); sham and contralateral tissue was also collected in regions matching the ipsilateral regions. To extract protein, all samples were homogenized in ice-cold lysis buffer (20mM Tris, 150mM NaCl, 0.05% Tween) and 1% each of 1 mmol/l PMSF (Sigma-Aldrich, P7626), 200mM Na3VO4 (Sigma-Aldrich, S6508) and a protease inhibitor cocktail (Sigma-Aldrich, P8340) followed by 2-h digestion at 4°C. Supernatant was collected after 15-min at centrifugation (14K x G) and analyzed for protein concentration using Bicinchoninic Acid kit (Thermo Fisher Scientific, #PI208609, IL,).

To quantify IBA1 and TMEM119 expression in each sample, 15 μg of protein was separated on a 10% Criterion® TGX™ Precast Protein Gel (BioRad Laboratories, 56671034, Hercules, CA) and transferred to a nitrocellulose membrane using a Trans-Blot TurboTransfer System (Bio-Rad Laboratories). A Chameleon® Duo Pre-stained Protein Ladder (LI-COR, 928-60000, Lincoln, NE,) was used for size reference and all blots were scanned on an Odyssey CLx imaging system. Total protein was calculated for each sample following incubation, according to the manufacturer’s specifications, in Revert 700 Total Protein Stain (LI-COR, 926-11010). Nonspecific binding was blocked with Odyssey Blocking Buffer (LI-COR, 927-50000) for 1 h at room temperature. The membrane was then cut between the 30Kda and 25Kda ladder (Chameleon Duo, 928-60000) and the top membrane with the larger proteins was incubated with rabbit anti-TMEM119 (1:500, Proteintech, 66948-1, IL) while the bottom portion was incubated with rabbit anti-IBA1 antibody (1:1000, Wako, 016-20001). Primary incubation was overnight at 4°C with a solution of LI-COR Odyssey Blocking Buffer and 0.2% Tween. Each membrane was washed (TBS-T and 0.2 Tween) and then incubated with IRDye® 800 CW goat anti-rabbit secondary antibody (1:10,000, LI-COR, 925-3221) diluted in Odyssey Blocking Buffer and 0.2% Tween for 1 h at room temperature. Total protein and bands at ∼45 KDa and ∼17KDa for TMEM119 and IBA1, respectively, were analyzed with Empiria Studio Software according to the manufacturer’s recommendations. All densitometry measurements were normalized to total protein prior to statistical analysis. All analysis was carried out by researchers blinded to the sex group.

Example of positive and negative control for TMEM119 were run with same conditions, but on a 10% Mini-Protean® TGX™ Precast Protein Gel (**Supplemental Figure 1**, BioRad, 4561036). Because of its limited use, a western analysis was used to test TMEM119 for its specificity using 3 µg of protein extracted from SH-SY5Y cells, mouse brain cortex homogenate, and mouse spleen homogenate. The SH-SY5Y cells (ATCC, VA) are a commonly used human derived neuroblastoma cell line and were the particular the cell line depicted by manufacturer https://www.ptglab.com/products/TMEM119-Antibody-27585-1-AP.htm). In this case, TMEM119 should be present in the SH-SY5Y and brain sample but not in the spleen. On the other hand, IBA1 should be present in all samples. In **Supplemental Figure 1** we show the expected TMEM119 band at ∼47kDa in the SH-SY5Y lane. Also, we show that while TMEM119 is present in the SH-SY5Y and brain samples, it is absent in the spleen sample. On the other hand, IBA1 is present in all three samples, SH-SY5Y, spleen, and brain samples.

### Statistical Analysis

Data are shown as mean ± SEM and were analyzed using GraphPad Prism 6.0 software (GraphPad Software, La Jolla, CA). Differences in infarct size, morphology, phagocytosis, TMEM119 immunohistochemistry and western blot were analyzed using non-repeated measure two-way analysis of variance (ANOVA) followed by a two-tailed Sidak multiple comparisons tests, or one-way ANOVA with Bonferroni *post hoc* test as appropriate.

## RESULTS

Representative images (**Figure 1A**) and summary data (**Figure 1B**) illustrate the distribution of brain infarct in the right hemisphere 60 min post-ischemia and 24 h after reperfusion between sex groups. Here, we show that infarct size is increased in male and post-menopause female mice versus pre-menopause mice (repeated measure two-way ANOVA: Section: F_(3, 93)_ = 22.5, p < 0.0001; Sex: F_(2, 31)_ = 4.9, p < 0.01; Interaction F_(6, 93)_ = 1.9, p < 0.05). Differences in infarct size between sex groups occur in the rostral brain regions (6 mm and 8 mm), but not caudal. All *post-hoc* analyses are reported in **Figure 1B**. We show that VCD, delivered *i*.*p*. when mice are 5-8 weeks of age, does not have an effect on brain infarct size after an ischemic stroke when delivered at 16 weeks of animal age (**Supplemental Figure 2**, repeated measure two-way ANOVA: Section: F_(3, 15)_ = 13.04, p < 0.001; VCD treatment: F_(1, 5)_ = 2.59, p > 0.05; Interaction: F_(3, 15)_ = 0.12, p > 0.05.).

**Figure 1.**
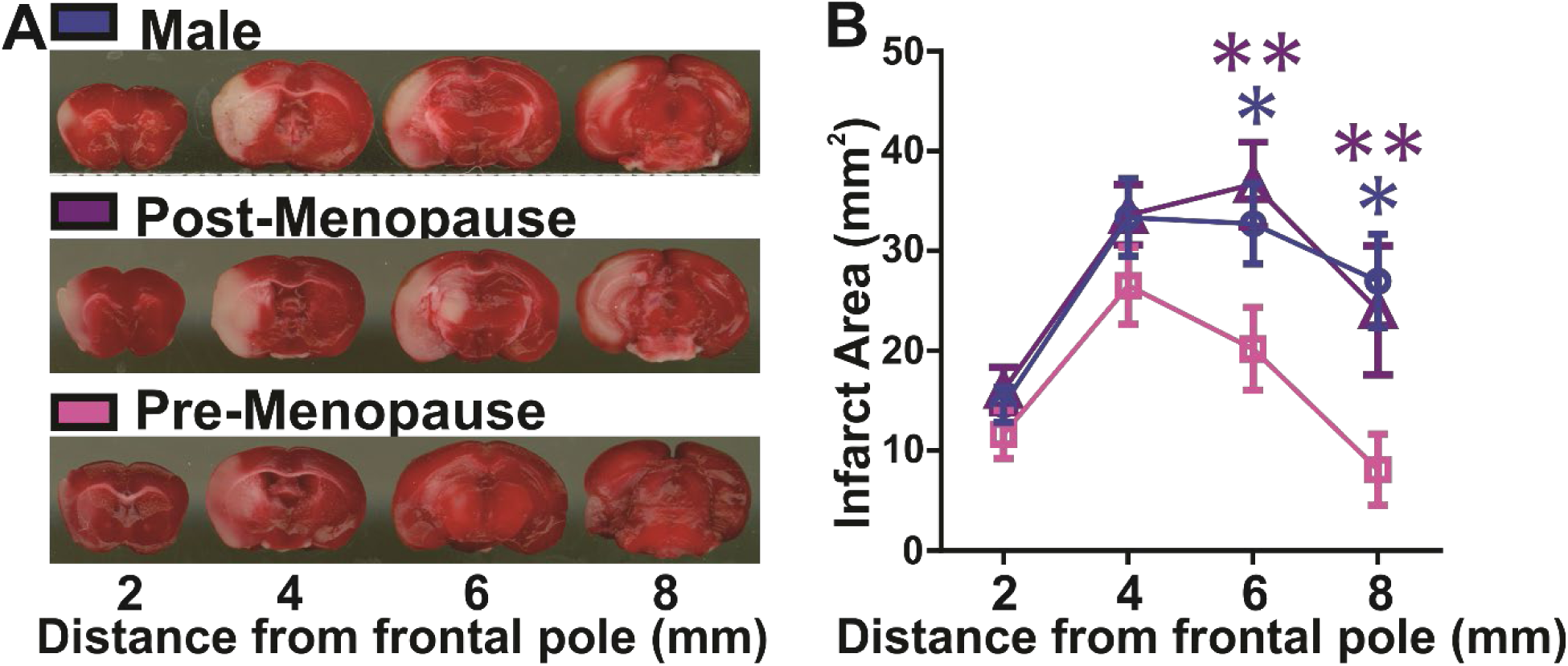
Brain infarct size is increased in male and post-menopausal mice versus pre-menopausal mice after ischemic stroke. **A)** Images of 2,3,5-triphenyltetrazolium chloride (TTC) stained brain sections in male and female post-menopause and pre-menopause mice after 60 min of ischemic stroke and 24 h of reperfusion. White area is necrotic and red area is healthy tissue. **B)** Summary data of infarct area (mm^2^) in a region between 2 mm-8 mm from frontal pole. Brain infarct area is increased in male or post-menopause mice versus pre-menopause mice. *, p < 0.05, ** p < 0.01. Male n = 13; Post-menopause female n = 11; Pre-menopause female n = 10

We next examined the contribution of sex and menopause on the morphologic and phagocytic responses of microglia to ischemic stroke. Total infarct volume was reduced in pre-menopause females as compared to male (p < 0.05) and post-menopausal female (p < 0.05) mice. However, at 4 mm from the frontal pole, infarct size was not significantly different across sex groups (one-way ANOVA, p = 0.3). Therefore, to avoid the confounding variable of infarct size differences among groups, all histological measures of microglial responses were assessed in coronal tissue sections 4 mm from the frontal pole, corresponding to Bregma 0 to +1. **Figure 2A** shows the brain regions imaged using a 40X objective for microglia morphology analysis. **Figure 2B** illustrates the IBA1 positive microglia imaged in each brain region imaged. We measured changes in microglia morphology among sex groups by quantifying the number of microglia endpoints and process length per cell via IHC and ImageJ analysis techniques. We show that the process endpoints on microglia decrease in regions that are proximal to the infarcted tissue similarly among sex groups (Region: F_(3, 45)_ = 88.45, p < 0.0001; Sex: F_(2, 15)_ = 2.26, p > 0.05; Interaction: F_(6, 45)_ = 1.12, p > 0.05). This effect is also observed for the summed process length per cell (Region: F_(3, 71)_ = 123.70, p < 0.0001; Sex: F_(2, 71)_ = 0.55, p > 0.05; Interaction: F_(6, 71)_ = 0.40, p > 0.05). All *post hoc* analyses are reported in **Figures 2C and 2D** (#, p < 0.0001 vs. sham region). To assess phagocytosis, we quantified the number of microglia with phagosomes, as identified by the ball and chain morphology shown in **Figure 3**. Our data show that microglia phagosomes are more present in the brain region that borders the necrotic tissue in all sex groups. However, sex and menopause influence microglia phagosomes differently in the distal and proximal brain regions (Region: F_(2, 45)_ = 44.54, p < 0.0001; Sex: F_(2, 45)_ = 3.31, p < 0.05; Interaction: F_(4, 45)_ = 4.22, p < 0.01). All *post-hoc* analyses are reported in **Figure 3** (**, p < 0.01 and #, p < 0.0001 vs. sham region; ^, p < 0.05 vs. male).

**Figure 2.**
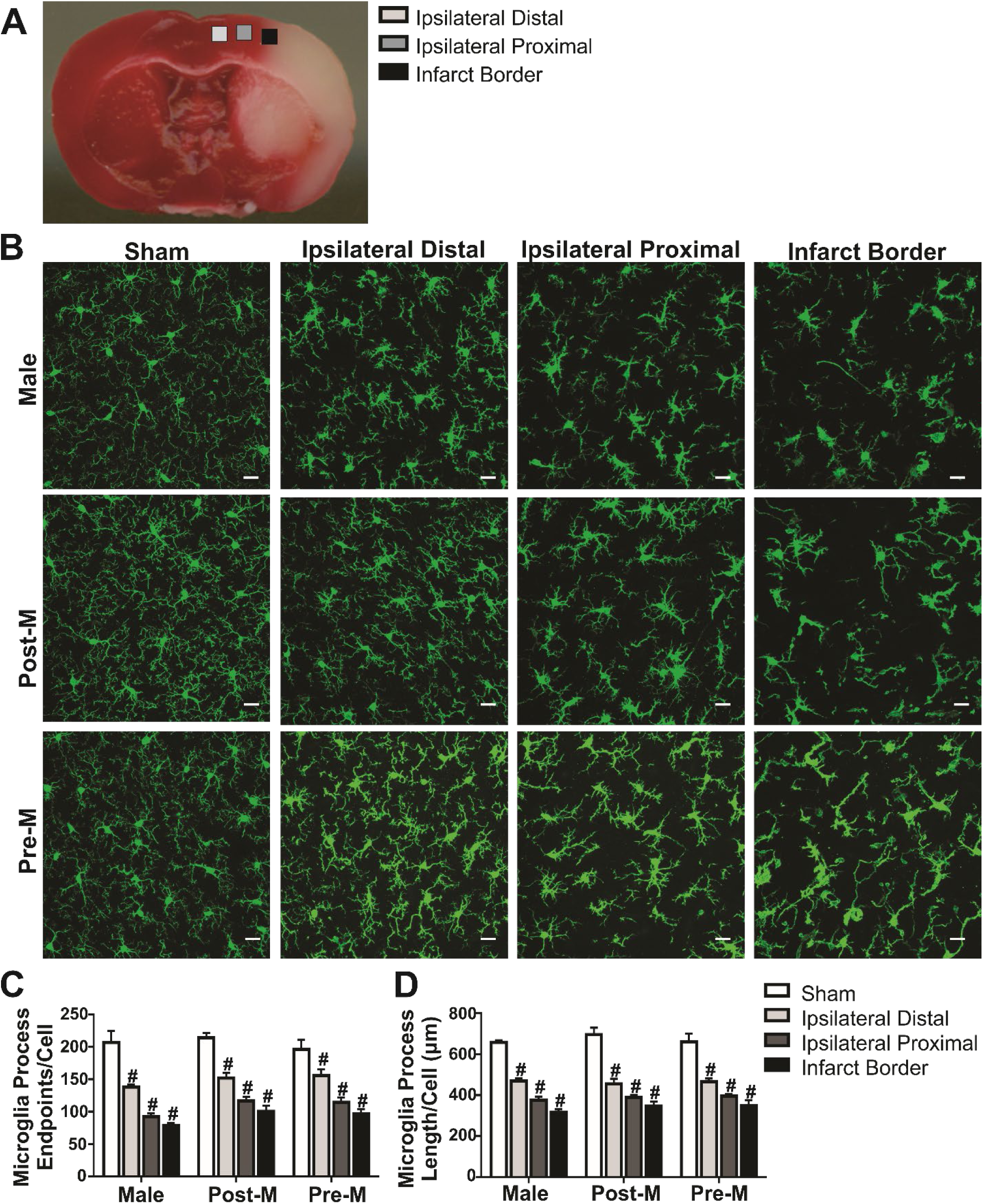
Microglia morphologic response to ischemic stroke is not influenced by sex or menopause. **A)** Images of IBA1 positive microglia (40X objective) in the cortex of mice after sham and stroke procedure. Three cortical ipsilateral regions in proximity to the infarcted tissue were included in the analysis as depicted in **(B)**. Summary data show that both the number of microglia process endpoints/cell **(C)** and summed process length/cell **(D)** decreases in proximity to the brain injury but with no sex differences. *Post-hoc* analysis is shown in the figure (****, p < 0.0001). Male n = 6; Post-menopause female n = 6; Pre-menopause female n = 6 for all regions. Scale bar = 20 µm.

We measured microglia TMEM119 immunofluorescence to assess if a portion of the IBA1 positive cells present in the ipsilateral brain regions after ischemic stroke were infiltrating macrophages. Our interest was specific to the infarct border region because of the decreased ramification and high level of phagosomes that may be indicative of a population of infiltrating macrophages rather than resident microglia. **Figure 4A** shows the brain regions imaged using a 20X objective to illustrate TMEM119 and IBA1 immunofluorescence with representative images shown in **Figure 4B**; cropped and merged images to show detail and co-localization of the two channels. Although the percent area of IBA1 immunofluorescence per frame is similar among brain regions and sex groups (Region: F_(2, 61)_ = 0.43, p > 0.05; Sex: F_(2, 61)_ = 3.57, p < 0.05 with no *post-hoc* differences; Interaction: F_(4, 61)_ = 1.23, p > 0.05), TMEM119 immunofluorescence is significantly decreased in the ipsilateral hemisphere proximal brain region with a similar effect observed among sex groups (Region: F_(2, 62)_ = 23.51, p < 0.0001; Sex: F_(2, 62)_ = 1.07, p > 0.05; Interaction: F_(4, 62)_ = 0.40; p > 0.05). All *post hoc* results are reported in Figures 4C and 4D (*, p < 0.05, **, p < 0.01 vs. sham region). Arrows in the cropped images in the ipsilateral proximal region highlight the observation that, although decreased, much of the remaining TMEM119 immunofluorescence was colocalized to phagosome-type structures (*i*.*e*. microglia processes with a ball and chain morphology).

**Figure 3.**
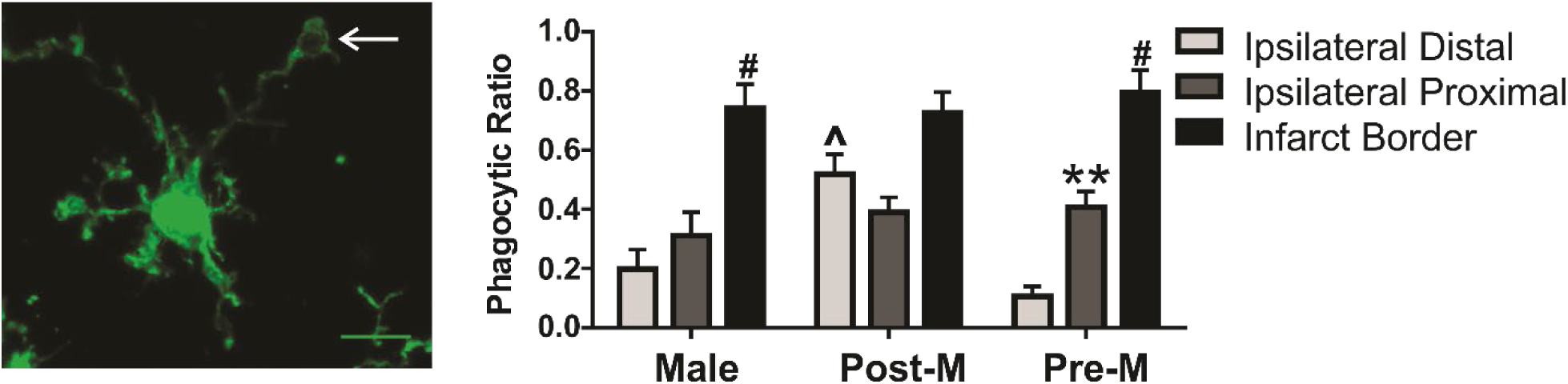
Microglia phagocytic responses are influenced by proximity to ischemic stroke, sex, and menopause. An example of microglia phagosome morphology (white arrow) and summary data of microglia phagocytic ratio (phagocytic cells/total microglia cells in the image frame) in ipsilateral hemisphere regions after ischemic stroke. Phagocytic ratio is increased in proximity to the infarcted region and is different according to sex and menopause. Regional *post-hoc* analyses (vs. ipsilateral distal region **, p < 0.01, ****, p < 0.0001) and sex group (^, p < 0.01 vs. male mice and p < 0.001 vs. pre-menopause female mice) are reported in the figure. Male n = 6; Post-menopause female n = 6; Pre-menopause female n = 6

**Figure 4.**
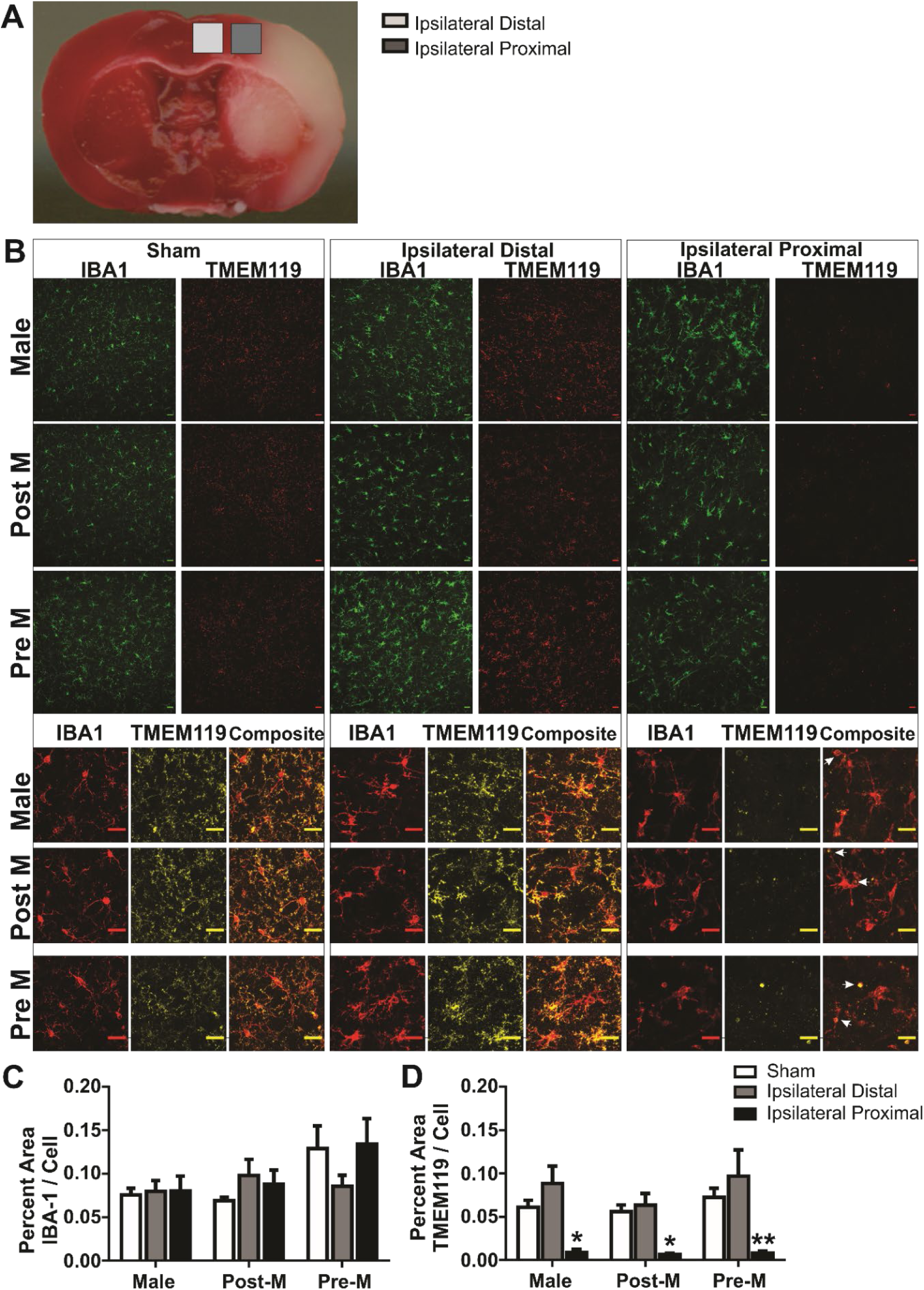
Percent area of TMEM119 immunofluorescence is decreased in proximity to the infarcted tissue. **A)** Images of IBA1 and TMEM119 immunofluorescence (20X objective) in the cortex of mice after sham and stroke procedure. Two brain regions (distal and proximal to injury) were imaged in the ipsilateral hemisphere. Below, cropped images show detail and composite show co-localization of IBA1 and TMEM119 immunofluorescence. **B)** Illustration of brain regions imaged. **C)** The percent area of IBA1 immunofluorescence per cells in frame remains unchanged among brain regions and sex. **D)** The percent area of TMEM119 immunofluorescence per IBA1 cells in frame is decreased in the ipsilateral proximal region versus sham in all sex groups. *, p < 0.05; **, p < 0.01. n = 6-9 in sex groups and regions. Scale bar = 20µm.

We further validated our immunofluorescence findings by measuring TMEM119 protein expression using western analysis. Brain tissue punches were carefully extracted from tissue regions similar to immunofluorescence regions as shown in **Figure 4A. Figures 5A-5C** are example blots of total protein, TMEM119 and IBA1, respectively. IBA1 protein expression was similar among brain regions and sex groups (**Figure 5D**; Region: Region: F_(3, 37)_ = 0.97, p > 0.05; Sex: F_(2, 37)_ = 2.38, p > 0.05; Interaction: F_(6, 37)_ = 0.73, p > 0.05). Different than IHC findings, summary data illustrate that TMEM119 protein levels are also similar among brain regions and sex groups (Region: F (3, 37) = 0.61, p > 0.05; Sex: F (2, 37) = 0.66, p > 0.05; Interaction: F (6, 37) = 0.61, p > 0.05). All *post hoc* results are reported in **Figures 5D and 5E**.

**Figure 5.**
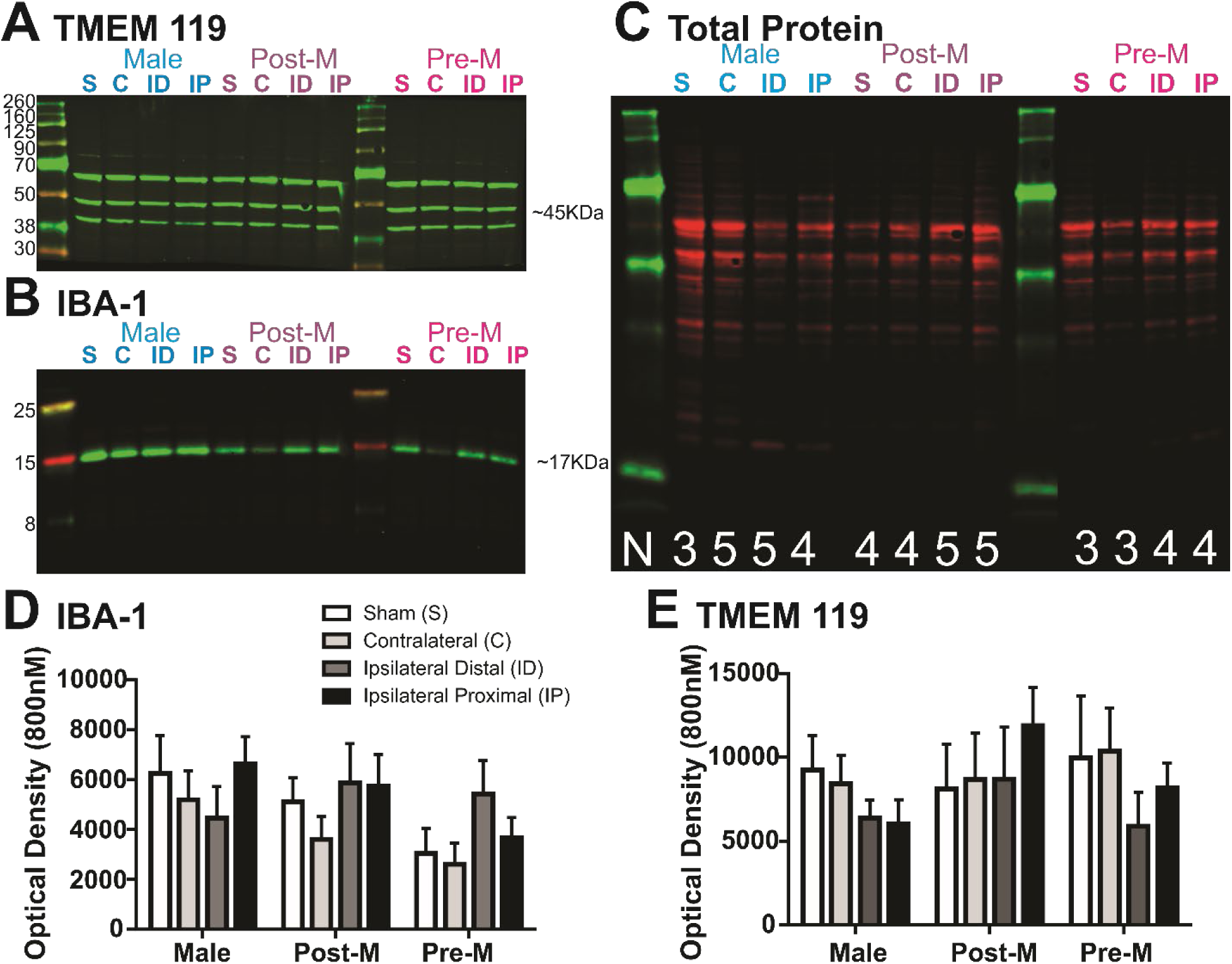
TMEM119 protein expression is unchanged in proximity to the infarcted tissue. Example blots of TMEM119 **(A)**, IBA1 **(B)**, and total protein **(C)** for regions and sex groups. **D)** Summary of IBA1 **(D)** and TMEM119 **(E)** data illustrate that these proteins remain relatively unchanged among brain regions and sex groups after ischemic stroke. Sample sizes range from 3-5 among sex groups and regions and are shown in **(A)**.

## DISCUSSION

Our primary purpose was to utilize both morphological analysis and a combination of TMEM119 immunofluorescence and protein expression to distinguish between microglial and infiltrating macrophage populations after ischemic stroke and 24 h of reperfusion. Our secondary purpose was to test for sex differences and, in females, the effect of menopause on microglia/macrophage cell populations after ischemic stroke. We are among the first to apply a post-menopause model that retains intact but follicle-depleted [36] ovaries to study sex differences in ischemic stroke injury and microglial responses. Using this model, we show that stroke infarct size is increased in male and postmenopausal female mice compared to premenopausal mice; this difference primarily occurs in caudal brain regions. The primary findings of this study are that while microglia become de-ramified in proximity to ischemic stroke, a large portion, if not all, of the imaged cells retained distinct cell processes. In addition, the ramified cells abutting necrotic tissue in the infarct border appear highly phagocytic as evidenced by prominent ball and chain morphologies. The lack of TMEM119 immunofluorescent-positive cells covered an unexpectedly large region that extended beyond the infarct border where infiltrating macrophages were most likely to have been present. Western blot analysis was used to validate the IHC findings. However, we show for the first time that the pattern of TMEM119 expression was not consistent between these methods. In light of these morphological and methodological incongruities, we are unable to definitively distinguish microglia from infiltrating macrophage cell populations in proximity to brain injury 24 h after ischemic stroke using TMEM119 alone. Instead, we suggest that in the context of acute ischemic stroke injury, TMEM119 is not a stable marker to denote microglia from infiltrating macrophages. The secondary findings of this study are that, when controlling for brain injury area, the transition of ramified microglia to less ramified cells was similar among male, pre- and postmenopause sex groups. We show evidence that phagocytic cells—cells with high phagocytic ratios—are distributed across a larger area in postmenopause mice than the male or premenopause groups in this preclinical model. Lastly, TMEM119 expression, whether measured via IHC or western blot methods, was similar among sex groups.

By combining a morphological assessment and a unique microglia marker, such as TMEM119, distinguishing between microglia and infiltrating macrophage populations might be possible in the healthy brain [25]. However, this approach was applied here to an acute brain injury model with gross brain injury, with inconclusive results. Microglia are known to be consistently distributed among the parenchyma with regions demarcated by their non-overlapping ramified morphology [39]. This pattern was observed in the present data with one exception—the region that directly abuts the infarct. In the infarct border region microglia distribution in the parenchyma becomes somewhat irregular as de-ramified cells were noted to be elongated and in possession of a great many phagosomes when compared to cells in other regions. Current literature suggests that the diapedesis of systemic cells into the parenchyma remain largely localized to the infarct core with little presence in the surrounding tissue at 24 h post-stroke [40]. Therefore, we hypothesized that cells absent of TMEM119 immunofluorescence would be more or less confined to the infarct border region. We found that cells with reduced TMEM119 immunofluorescence area extended well into the proximal, and almost to the distal region. In these regions cells possessed an abundance of processes and were uniformly positioned in the parenchyma—both characteristics of microglia [25]. This led us to verify our IHC findings using western blot methods, which revealed a discrepancy. A few scenarios may account for the conflicting results between methods. First, it is often the case that antibodies successfully used in immunohistochemistry methods are not always successful in western blot methods due to sample preparations. This was the case and a limitation of this study; two different antibodies were assessed, and neither worked in the complementary method. However, we included appropriate positive and negative controls to ensure the validity of these manufacturer prepared antibodies.

Second, it is possible that TMEM119 possesses a functional response to a gross injury such as ischemic stroke. In fact, TMEM119 was not entirely absent from ramified cells in the proximal region and, when present, appeared to colocalize to phagosomes. This observation may suggest a yet undetermined response to injury associated with phagocytosis. Others have demonstrated that, in cultured human microglia, *TMEM19* transcripts are reduced by the presence of the interleukin IL-4 [41] or interferon gamma (INFγ), but not LPS [29], which could indicate an injury specific transcriptional change.

Transmembrane proteins are part of a large family that, in general, consist of anchored proteins thought to act as transport channels across the lipid bilayer [27, 28]. However, by and large, the exact functions of these abundant proteins, including TMEM119 [30], remains unknown [28]. TMEM119 was originally identified as an osteoblast inhibitory factor (OBIF) derived from osteoclast cell lines [42, 43]. Notably, osteoclasts are cells derived from the monocyte/macrophage cell line [44]. *Obif* mRNA is present in a diversity of tissues (https://www.ncbi.nlm.nih.gov/gene/231633), and while Mizuhashi et al. (2015) describes bone mineralization and spermatogenesis phenotypes in *Obif*^−/−^ mice, a central nervous system phenotype was not investigated [45]. Others have demonstrated that TMEM119 has an important function in the differentiation of myoblasts into osteoblasts [29, 46-50]. Investigations of TMEM119 in the brain centers around novel methodological tools to distinguish microglia from other myeloid populations [25, 51] and descriptions of cell populations in health and disease [29-31, 52, 53]. Despite these recent increases in utilizing TMEM119 in brain research, its function in the brain remains unknown. In our images, TMEM119 was rarely completely absent from IBA1 positive cells but rather, the area of immunofluorescence was visibly diminished and where present, often co-localized to microglia phagosomes or processes. This observation suggests to us that either the TMEM119 epitope used for antibody-antigen interactions in IHC preparations may be degraded with injury or that this transmembrane protein has a functional response to injury that must be further characterized. Similar to another microglia distinguishing protein purinergic receptor P2Y12 [54], TMEM119 expression may change according to physiologic or pathologic stimuli or to the local inflammatory environment. Therefore, its methodological application may be most useful in discriminating among populations of microglial responses to injury rather than distinguishing among myeloid populations.

It is well established that sex is an important biological variable in determining ischemic stroke outcomes [3, 55]. However, the mechanisms that underlie observed sex differences in brain injury outcomes are not fully established. Therefore, we included the three relevant sex groups, pre- and postmenopause females as well as males. We posited that differences in cell populations (microglia vs. infiltrating macrophages) among these sex groups would be insightful to understand disparate injury evolution and stroke outcomes. We used a postmenopause model that induced ovarian failure rather than ovariectomy to best represent the human population, where surgically induced menopause is less common than naturally occurring menopause. Using this model, we replicated previous research, primarily carried out in an ovariectomized rodent models, that male and postmenopause female mice have more severe stroke injury when compared to premenopausal female mice [3]. However, infarct size was similar between age-matched postmenopause female mice and male mice. By using a limited region of interest for our investigation, we were able control for infarct size in the immediate region of our investigation. Our findings suggest that sex and menopause are not factors that influence the morphological response of microglia to ischemic stroke injury or post-stroke microglia TMEM119 protein expression. Considering the similarity in findings across groups, whether it be microglia or macrophages, cell responders appear to be similar regardless of sex or menopause status.

## CONCLUSIONS

We demonstrate that the accelerated ovarian failure model, wherein mouse ovaries remain intact but dysfunctional, can be utilized to investigate ischemic stroke pathophysiology. Using this model, we illustrate that microglia transition into less ramified cells when in proximity to the infarct but without a sex or menopause effect. On the other hand, the phagocytic nature of microglia indicated by morphology, is increased in regions that are distal to the infarcted region in postmenopause mice when compared to the other sex groups and may contribute to disparities in post-stroke injury. This is an area for future investigation. While TMEM119 remains a promising addition to the repertoire of markers used to study microglia, the results of this study demonstrate that it cannot be used to distinguish between resident microglia and infiltrating macrophages in all cases. The disparity between IHC and Western analysis suggests that the function of TMEM119 is more nuanced than initially thought, and that its efficacy as a reliable microglial marker may be both function- and response-specific. Furthermore, the apparent localization of TMEM119 immunofluorescence to microglial processes, and particularly to phagosomes when present in proximity to injury, suggests that its function may be specific to filopodia extension, cell mobility, and an actively phagocytic microglial response.

## LIST OF ABBREVIATIONS

TMEM119: transmembrane protein 119
VCD: 4-vinylcyclohexene diepoxide
MCA: middle cerebral artery
IBA1: ionized calcium binding adaptor molecule 1
ANOVA: analysis of variance
IHC: immunohistochemistry
IL-4: interleukin 4
INFγ: interferon gamma
LPS: lipopolysaccharide
OBIF: osteoblast inhibitory factor

## DECLARATIONS

Ethics approval and consent to participate

All animal handling and experiments were performed according to methods approved by and in compliance with the University of Arizona Institutional Animal Care and Use Committee and according to the National Institutes of Health guide for the care and use of laboratory animals (protocol approval number 14-539).

## Consent for publication

Not applicable

## Availability of data and materials

The datasets used during the current study are available from the corresponding author on reasonable request.

## Authors’ contributions

KFY assisted in conceptualizing the study, carried out microglia analysis blinded to the experimental groups, and edited the manuscript. RG assisted with western blot experiments and analysis blinded to the experimental group. VCS and SAW assisted with microglial analysis blinded to experimental group. MJB and TF assisted with western blot experiments, blinded analysis, and provided manuscript editing. HWM conceptualized the study, conducted stroke surgeries, carried out all experiments and image acquisition, and wrote the manuscript.

## Acknowledgements

I would like to acknowledge the Microscopy Alliance at the University of Arizona and the expertise of Patricia Jansma, MS for her assistance with image acquisition on the Marley Zeiss LSM880 NLO upright multiphoton/confocal microscope. Internal funding from the University of Arizona College of Nursing Emmons Award and Bauwens Award funded this research.

---

**Supplemental Figure 1.**
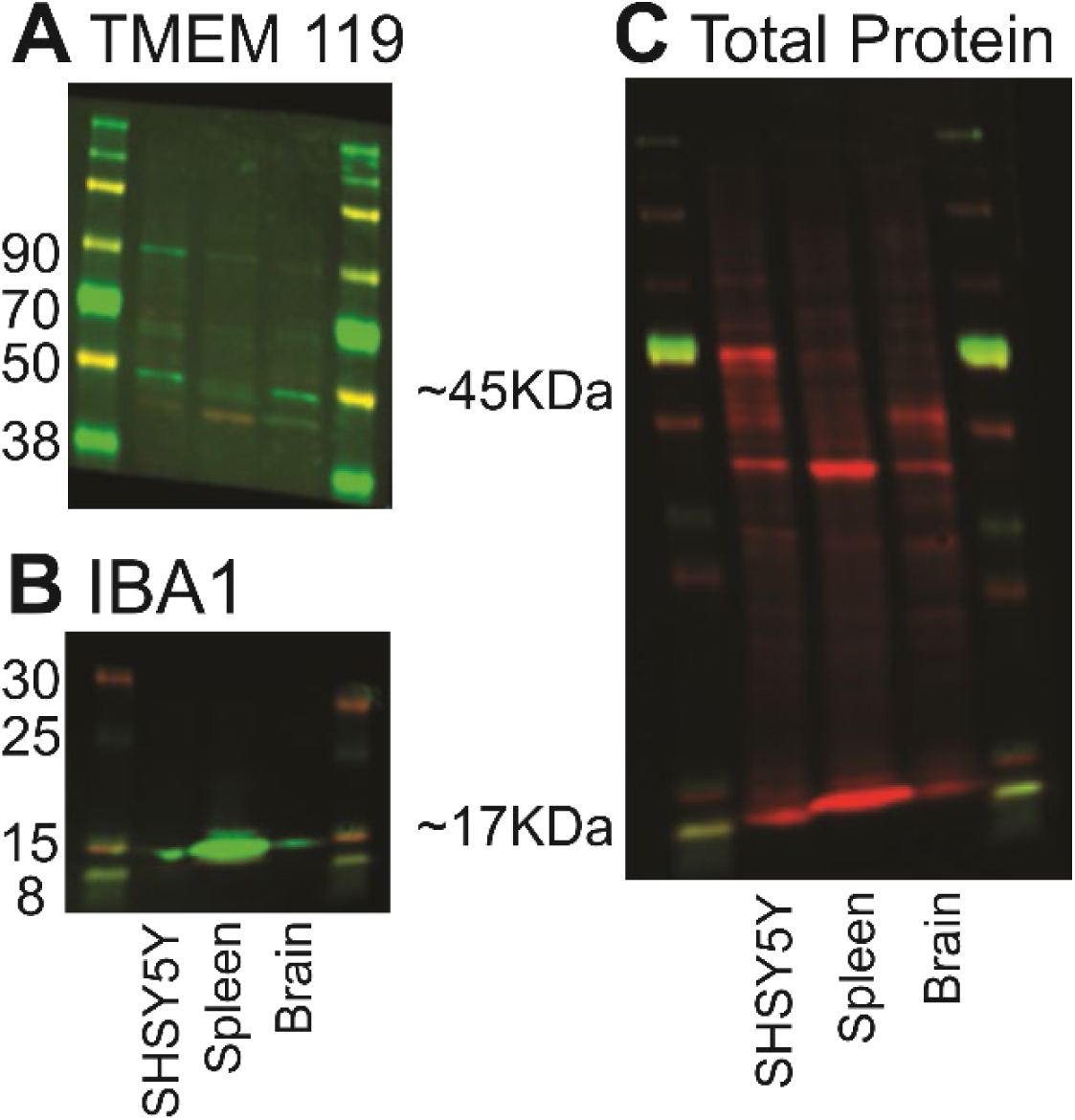
TMEM119 antibody validation in control samples. Example blots of TMEM119 **(A)**, IBA1 **(B)**, and total protein **(C)** in SH-SY5Y cells (lane 1), spleen (lane 2) and brain (lane 3) samples. This image illustrates that while TMEM119 is present in the SH-SY5Y cells (tested by manufacturer) and brain samples, it is not present in the spleen. IBA1 is present in all samples.

**Supplemental Figure 2.**
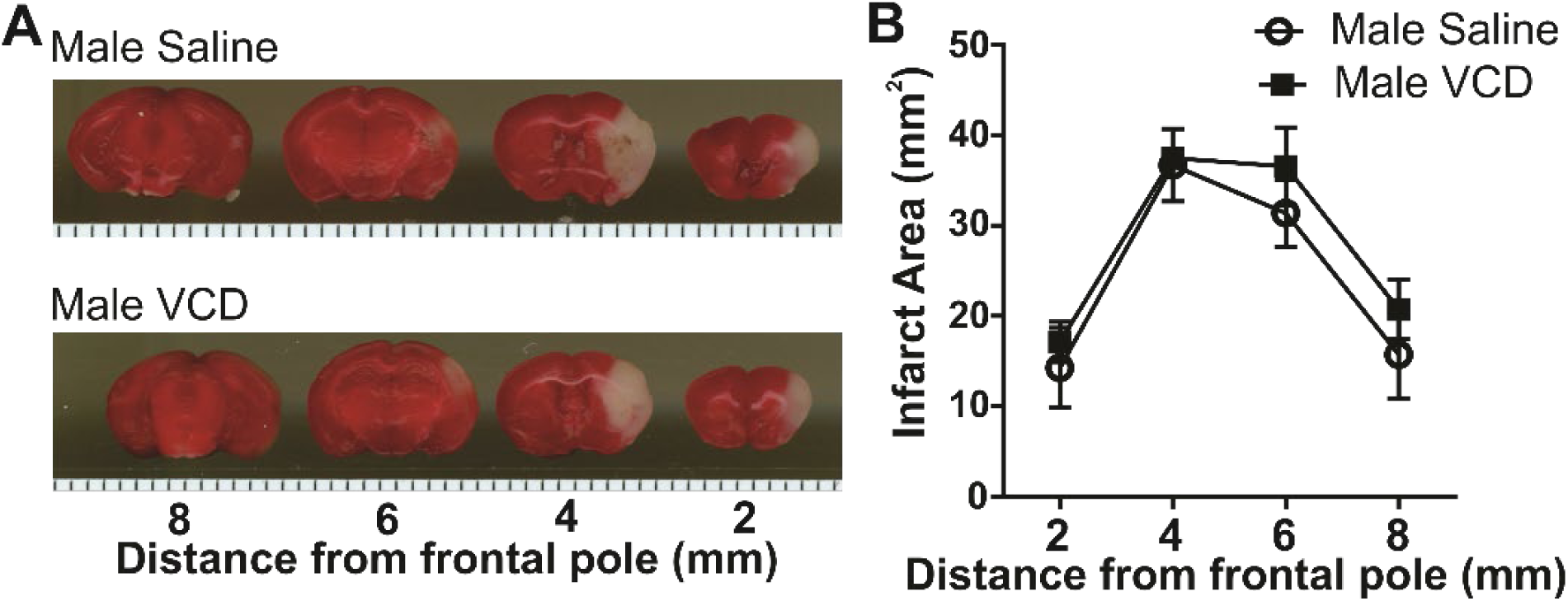
VCD injections do not affect stroke size in male mice. **A)** Images of 2,3,5-triphenyltetrazolium chloride stained brain sections in male mice treated with saline and male mice treated with VCD after 60 min of ischemic stroke and 24 h of reperfusion. White area is necrotic and red area is healthy tissue. **B)** Summary data of infarct area (mm^2^) between 2 mm-8 mm from frontal pole. Brain infarct area is not significantly different between saline and VCD treated mice. Sample size: Male saline n = 4, Male VCD n = 3.

